# Neutrophil extracellular traps arm DC vaccination against NPM-mutant myeloproliferation

**DOI:** 10.1101/2021.05.19.444838

**Authors:** Claudio Tripodo, Barbara Bassani, Elena Jachetti, Valeria Cancila, Claudia Chiodoni, Paola Portararo, Laura Botti, Cesare Valenti, Milena Perrone, Maurilio Ponzoni, Patrizia Comoli, Antonio Curti, Mario Paolo Colombo, Sabina Sangaletti

## Abstract

Neutrophil extracellular traps (NET) are web-like chromatin structures composed by dsDNA and histones, decorated with anti-microbial proteins. Their interaction with dendritic cells (DC) allows DC activation and maturation toward presentation of NET-associated antigens. Differently from other types of cell death that imply protein denaturation, NETosis preserves the proteins localized onto the DNA threads for proper enzymatic activity and conformational status, including immunogenic epitopes. Besides neutrophils, leukemic cells can release extracellular traps displaying leukemia-associated antigens, prototypically mutant nucleophosmin (NPMc+) that upon mutation translocates from nucleolus to the cytoplasm localizing onto NET threads. We tested NPMc+ immunogenicity through a NET/DC vaccine to treat NPMc-driven myeloproliferation in transgenic and transplantable models. Vaccination with DC loaded with NPMc+ NET (NPMc+ NET/DC) reduced myeloproliferation in transgenic mice, favoring the development of antibodies to mutant NPMc and and the induction of a CD8+ T cell response. The efficacy of this vaccine was also tested in mixed NPMc/WT bone marrow chimeras in a competitive bone marrow transplantation setting, where the NPMc+ NET/DC vaccination impaired the expansion of NPMc+ in favor of WT myeloid compartment. NPMc+ NET/DC vaccination also achieved control of an aggressive leukemia transduced with mutant NPMc, effectively inducing an anti-leukemia CD8 T cell memory response.

**eTOC Summary:** *NPMc is an AML-triggering mutation inducing translocation of NPM into the cytoplasm, allowing its association to NET threads. On NET, NPMc becomes immunogenic acting as an alarmin and exposing immunogenic peptides. We demonstrate that NET can be exploited as anti-tumor weapons in DC-based vaccination strategies*.

## Introduction

Fifteen years ago, a scanning electron microscopy image from Volker Brinkmann showed for the first time spider web-like chromatin structures extruded by neutrophils, called neutrophil extracellular traps (NET), to entrap fungi and bacteria (1). Such extracellular chromatin is composed by DNA and histones also decorated with anti-microbial proteins like myeloperoxidase (MPO) and neutrophil elastase suggesting the hypothesis that eukaryotic chromatin evolved under the need of maintaining genome integrity while defending the organism (2). Extracellular traps can be released by other innate immune cells including eosinophils, macrophages and mast cells (3) and, as recently shown, by cells of the adaptive immunity like CD4 T-helper cells (4). Additionally, since their discovery, NET have been associated with inflammatory and immune-mediated diseases like diabetes, arthritis, systemic vasculitis and lupus erythematosus (5). In a previous study we demonstrated that the adoption of a NET-based DC vaccination was able to break tolerance against neutrophil cytoplasmatic antigens and induce anti-neutrophil cytoplasmatic antibodies (ANCA)-associated autoimmunity^6^. Indeed, NET are highly immunogenic by virtue of the immune adjuvant effect of DNA and its associated proteins, as well as for the sticky properties of the DNA thread that enable NET persistent interaction with dendritic cells (DC) for efficient loading of antigens followed by DC maturation and migration to draining lymph nodes for cross-presentation (6).

Also, myeloid transformed cells can extrude NET into the extracellular space to activate the contact system of coagulation (7) or to sustain myeloproliferation. We have recently described extracellular traps enrichment in bone marrow (BM) biopsies from *NPM1* mutant AML patients and, using an ad-hoc transgenic mouse, we showed that NPM cytoplasmic compartmentalization allows mutant NPM (NPMc+) to be re-localized onto the NET threads, exerting alarmin functions (8). The co-localization of histones with NPMc*+* is a supporting evidence of their direct release from the leukemic clone.

The immune system has the capacity to eradicate acute myeloid leukemia (AML) as shown by the graft-versus-leukemia effect that is obtained after allogeneic hematopoietic stem cell transplantation (HSCT). Accordingly, ideal immune molecular targets of AML should be restricted to leukemic cells, expressed by most of them (encompassing also leukemic stem cells), instrumental to maintain the leukemic status, immunogenic, and clinically effective.

Some leukemia-associated antigens described so far, such as, RHAMM, Proteinase 3, and Wilms’ tumor antigen-1 (WT-1) have entered clinical trials as peptide vaccines (9).

*NPM1* mutations are among the most frequent molecular alterations in AML, where they play a prognostic role (10). *NPM1* gene encodes for the nucleolar protein nucleophosmin that regulates the ARF-p53 tumor-suppressor pathway (11). NPMc mutations cause the stable relocalization of the protein from nucleus to cytoplasm, an event that, per se, is sufficient to trigger AML (12). The improved overall survival of patients with NPMc+ AML is has been possibly explained by a T-cell response against the mutant epitopes. The specific CD4^+^ and CD8^+^ T cell response against these epitopes raised the possibility of exploiting such property to immunize NPMc+ patients to control MRD or during maintenance treatment (13). In this context, NET directly released by leukemic cells could efficiently display tumor-specific associated antigens and be used as vehicles for new DC-based vaccines.

In this study we investigated whether the AML-associated NPMc is immunogenic whenever it is part of the NET threads and whether NPMc+ blast-derived NET could be adopted in vaccination strategies to control myeloproliferation or leukemia outgrowth in transgenic or transplantable mouse models.

## Results

### NPMc+ NET/DC immunization controls NPMc-driven myeloproliferation

h-MRP8-*NPM1+* (NPMc*+*) transgenic mice develop a myeloproliferation with expansion of mature CD11b+ myeloid cells and Gr-1+c-Kit+ myeloblasts, without development of overt acute leukemia. Using NET from NPMc*+* transgenic mice we previously demonstrated that mutant NPM can work as an alarmin when localizing in the cytoplasm and eventually transferred onto the NET thread^17^, becoming immunogenic. In this work we tested the possibility of using NPMc*+* NET in a DC-based vaccination strategy to control myeloproliferation of NPMc*+* transgenic mice. To this end, mice were immunized with DC co-cultured with NPMc*+* NET or WT NET (Figure 1A). During co-culture, DC became loaded with the prototypical NET-associated antigen MPO (Figure 1B, red signal) and, only in case of NPMc*+* NET, with mutant NPMc (Figure 1B, green signal). NPMc*+* NET/DC immunization, but not immunization with WT NET/DC, conspicuously reduced the signs of myeloid expansion in the BM of NPMc*+* mice, with a decrease in dense clusters of morphologically immature granulocytic elements, as shown by histopathological analysis of H&E-stained BM sections (Figure 1C), and a reduction in the presence of myeloid blasts in favor of more segmented forms on BM blood smears (Figure 1C). Moreover, NPMc*+* NET immunization resulted in a significant decrease of cytoplasmic NPM-expressing elements, as assessed by immunofluorescence (Figure 1D-E). Accordingly, FACS analysis of the PB confirmed the reduction of circulating immature GR-1+c-Kit+ precursors and of the overall frequency of CD11b+ cells (Figure 1F-G) in mice vaccinated with DC loaded with NPMc*+* NET.

**Figure 1.**
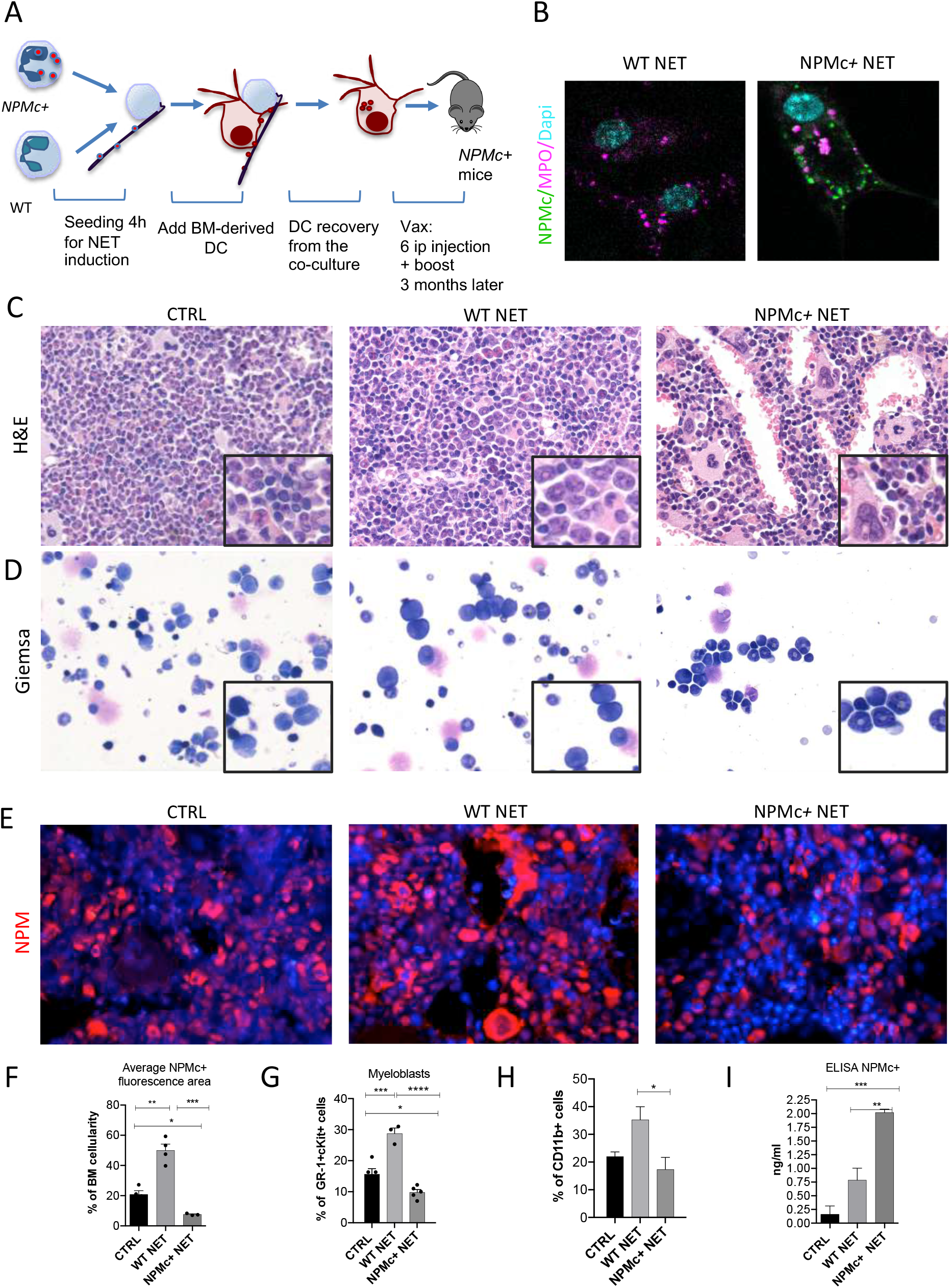
Vaccination with NPM+ NET/DC controls NPMc+-driven myeloproliferation. A. Schematic representation of the vaccination experiment. B. IF analysis for MPO (purple) and NPM (green) of DC co-cultured with NPMc*+* or WT NET; C. BM histopathology of NPMc*+* transgenic mice vaccinated with WT or NPMc+ NET-loaded DC or left untreated; D. May-Grunwald Giemsa staining of BM smears from NPMc*+* transgenic mice vaccinated with WT or NPMc+ NET-loaded DC or left untreated. E. IF analysis for NPM on BM sections from vaccinated or control mice; F. Quantification of NPMc*+* areas in the IF analysis; PB FACS analysis for GR-1+c-Kit+ myeloblasts (G) and CD11b+ myeloid cells (H) NPMc*+* transgenic mice vaccinated with WT or NPMc+ NET-loaded DC or left untreated. I. Quantification of auto-antibodies to mutant NPM developing in the serum of vaccinated mice. Statistical analysis (panel F-I):one-way ANOVA; Tukey’s multiple comparison test. * p< 0.05;**p<0.01;***p<0.001 (n=5/group).

The immune response triggered by NPMc*+* NET immunization induced anti-NPMc*+* serum antibodies detected by ELISA (Figure 1H) and the increase of CD8+ T-cells frequency in BM infiltrates (Figure 2A-B). Of note, the increased infiltration of CD8+ T-cells associated with a higher frequency of CD8+ T cells closely contacting NPMc*+* cells in the BM of vaccinated mice (Figure 2C-D). Overall, these data indicate that DC vaccination with NPMc+ NET is able to induce immune activation towards control of NPMc+ myeloproliferation.

**Figure 2.**
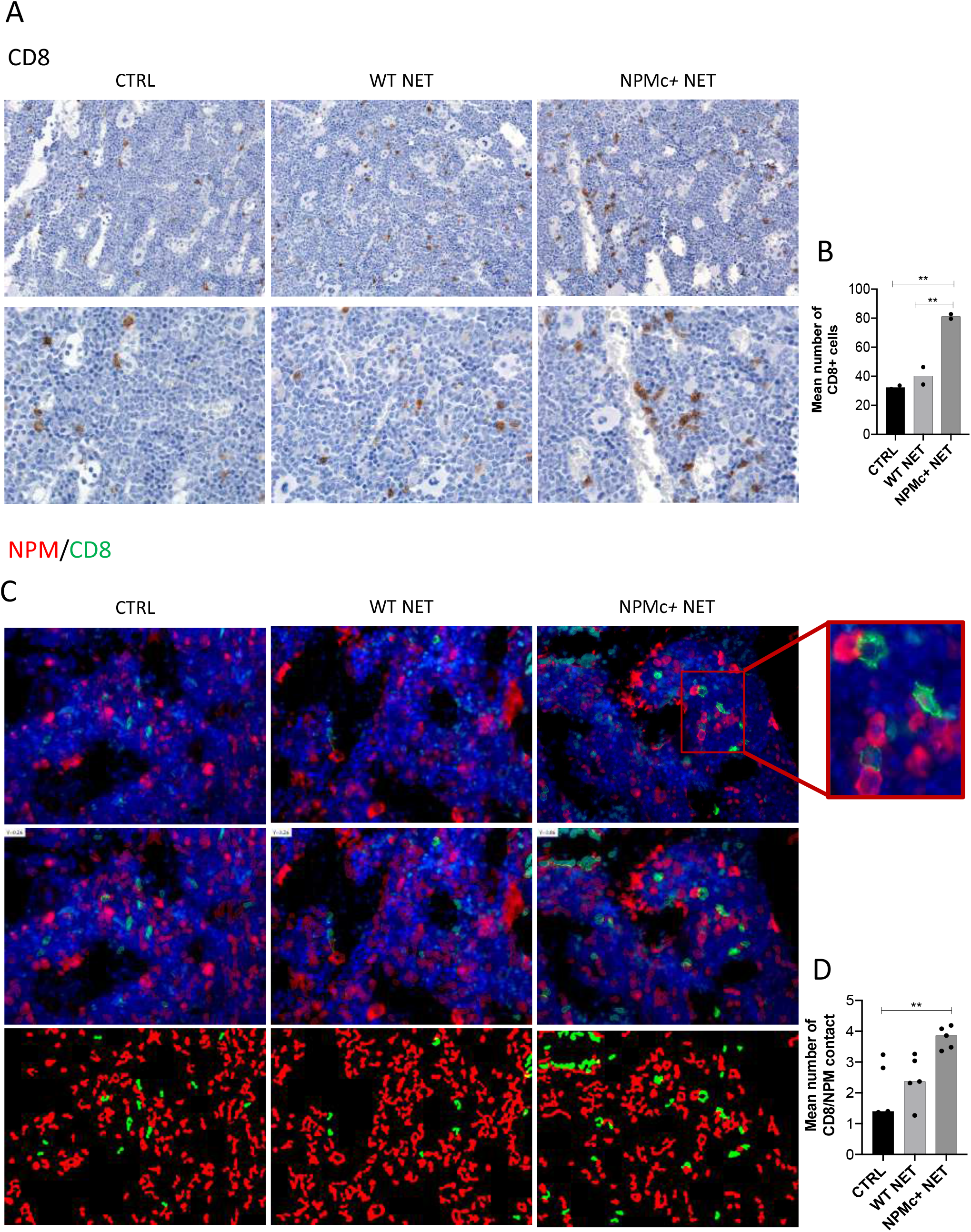
Analysis of CD8 T cell frequency and interaction with NPMc+ cells in BM biopsies from control and vaccinated mice. A. IHC analysis of CD8+ cells in BM sections of NPMc+ transgenic mice vaccinated with WT or NPMc+ NET-loaded DC or left untreated. B. Quantification of CD8+ T cells performed by counting the number of immunoreactive cells out of five non-overlapping high-power (x400) microscopic fields for every BM sample. C. Representative IF analysis on BM sections of NPMc+ transgenic mice vaccinated with WT or NPMc+ NET-loaded DC or left untreated, showing the reciprocal distribution of CD8+ T cells (green) and NPMc+ (red) B. Software-based quantitative analysis of cell-cell contact between CD8+ and NPMc+ cells on segmented IF microphotographs. Statistical analysis: one-way ANOVA; Tukey’s multiple comparison test. **p<0.001.

### NPMc+ NET/DC immunization selectively impairs NPMc+ mutant hematopoiesis in competitive BMT setting

To test the activity of the DC/NPMc*+* NET vaccination in controlling the expansion of NPMc*+* cells, we performed competitive bone marrow transplantation experiments in which WT mice were transplanted with a 1:1 mixture of Lin-precursors from NPMc*+* Tg mice (CD45.1) and WT mice (CD45.2). Four weeks after BMT mice were vaccinated with DC loaded with either WT or NPMc*+* NET (Figure 3A). Mice were sacrificed 12 weeks after BMT and analyzed for the frequency of circulating CD45.1 (NPMc+derived) and CD45.2 (WT-derived) CD11b+ and GR-1+c-Kit+ myeloblasts, respectively. Representative analysis (Figure 3B) and cumulative data (Figure 3C) show that vaccination with NPMc*+* NET-loaded DC impaired the expansion of CD45.1 mutant hematopoietic cells in favor of CD45.2 WT cells in the competitive setting. These results suggest that vaccination with NET carrying mutant NPMc could selectively control the expansion of mutant cells derived from NPMc*+* precursors, while sparing the normal counterpart. Interestingly, the reduced expansion of CD45.1 NPMc*+* cells was associated to increased CD8 T-cell infiltration in vaccinated mice (Figure 3D-E). The selective effect on mutant cells in the competitive setting underscores the efficacy and specificity of the engendered immune response towards mutant hematopoiesis.

**Figure 3.**
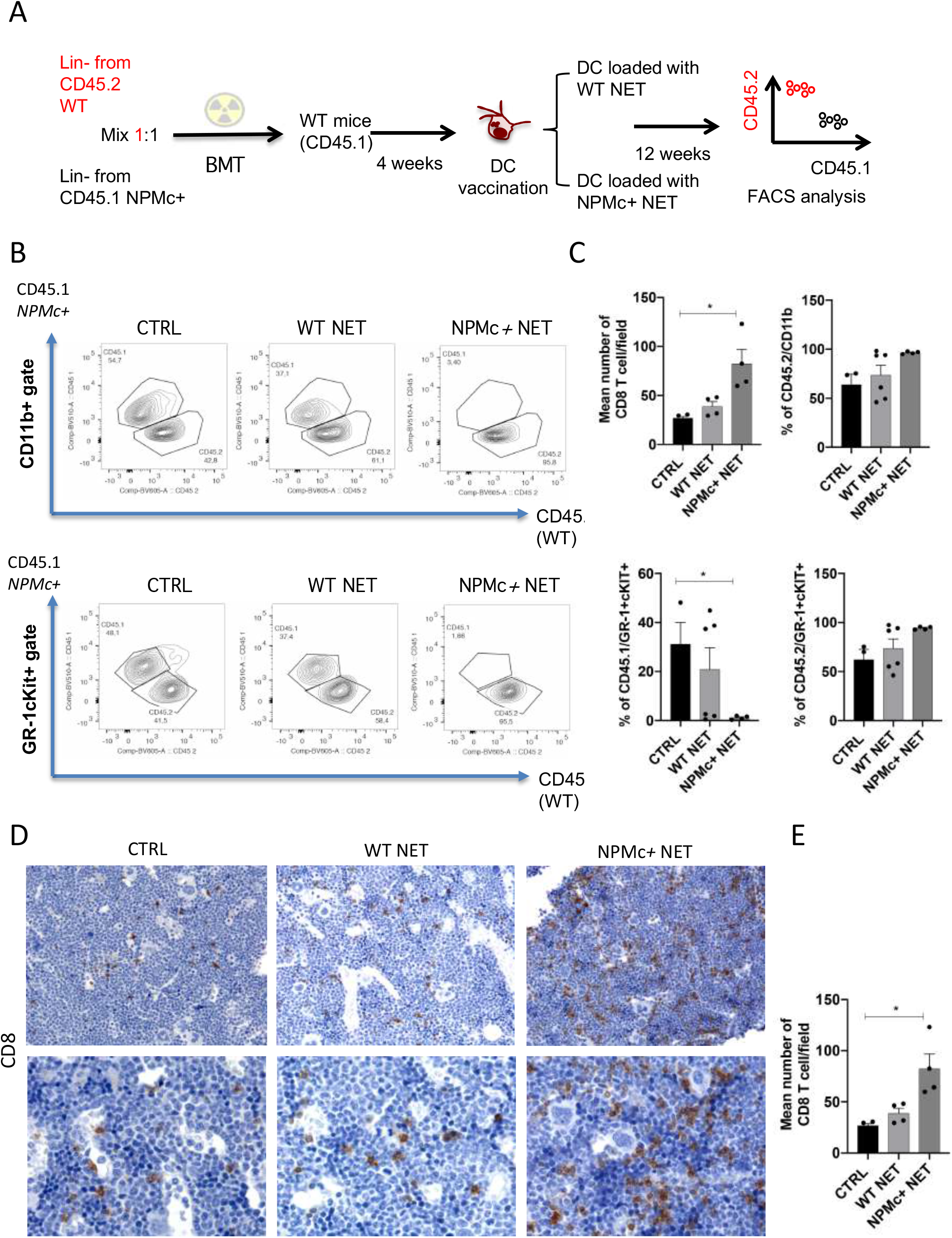
Vaccination with with NPM+ NET/DC controls the expansion of NPMc+ cells in competitive BMT assay. A. Schematic representation of the competitive BMT experiment; B. Representative dot plots showing the frequency of CD45.1 (NPMc+) and CD45.2 (WT) in myeloid cell- (CD11b+) and myeloblast- (GR-1+c-Kit+) gate of BM chimeras that received vaccination with WT or NPMc+ NET/DC; C. Cumulative data showing the frequency of CD45.1 (NPMc+) and CD45.2 (WT) within the CD11b+ and GR-1+c-Kit+ gate. Statistical analysis: Mann-Whitney test *p<0.05. (n=4/group, one representative experiment out of 3 performed). A. IHC analysis of CD8+ cells in BM sections of in BM chimeras vaccinated with WT or NPMc+ NET-loaded DC or left untreated. B. Quantification of CD8+ T cells performed by counting the number of immunoreactive cells out of five non-overlapping high-power (x400) microscopic fields for every BM sample. Statistical analysis: one-way ANOVA; Tukey’s multiple comparison test. **p<0.001.

### NPMc+ NET/DC vaccination prevents transplantable NPMc+ leukemia *cell growth*

Since the indolent phenotype of NPMc*+* Tg mice did not allow a short-term readout for assessing vaccine efficacy, we generated a transplantable leukemia model expressing mutant NPMc. To this purpose, the leukemia cell line C1498 was infected with a lentiviral vector expressing mutant *NPM1* (C1498-NPMc*+*) (Suppl. Figure 1A) and then injected into NPMc*+* Tg mice. Given that this injection in the transgenic mice is expected to induce tolerance in case of potential transgene antigenicity (14), we tested with this approach whether NPMc*+* NET-based vaccine could disrupt tolerance in NPMc*+* mice inducing an immune response able to control C1498-NPMc+ leukemia. In this experiment DC uploaded with NPMc*+* NET were injected intradermal in NPMc+ Tg mice bearing the C1498-NPMc*+* leukemia. The vaccine was administered at day 3, 5, 10 and 14 post leukemia injection (Figure 4A). Tumor growth was monitored twice a week and the raising of NPMc-specific CD8+ T cells was evaluated through *in vivo* cytotoxicity assay ^21^ by injecting mice with splenocytes pulsed 1 hour with NPMc*-*derived MHC-I peptides or with an unrelated peptide.

**Figure 4.**
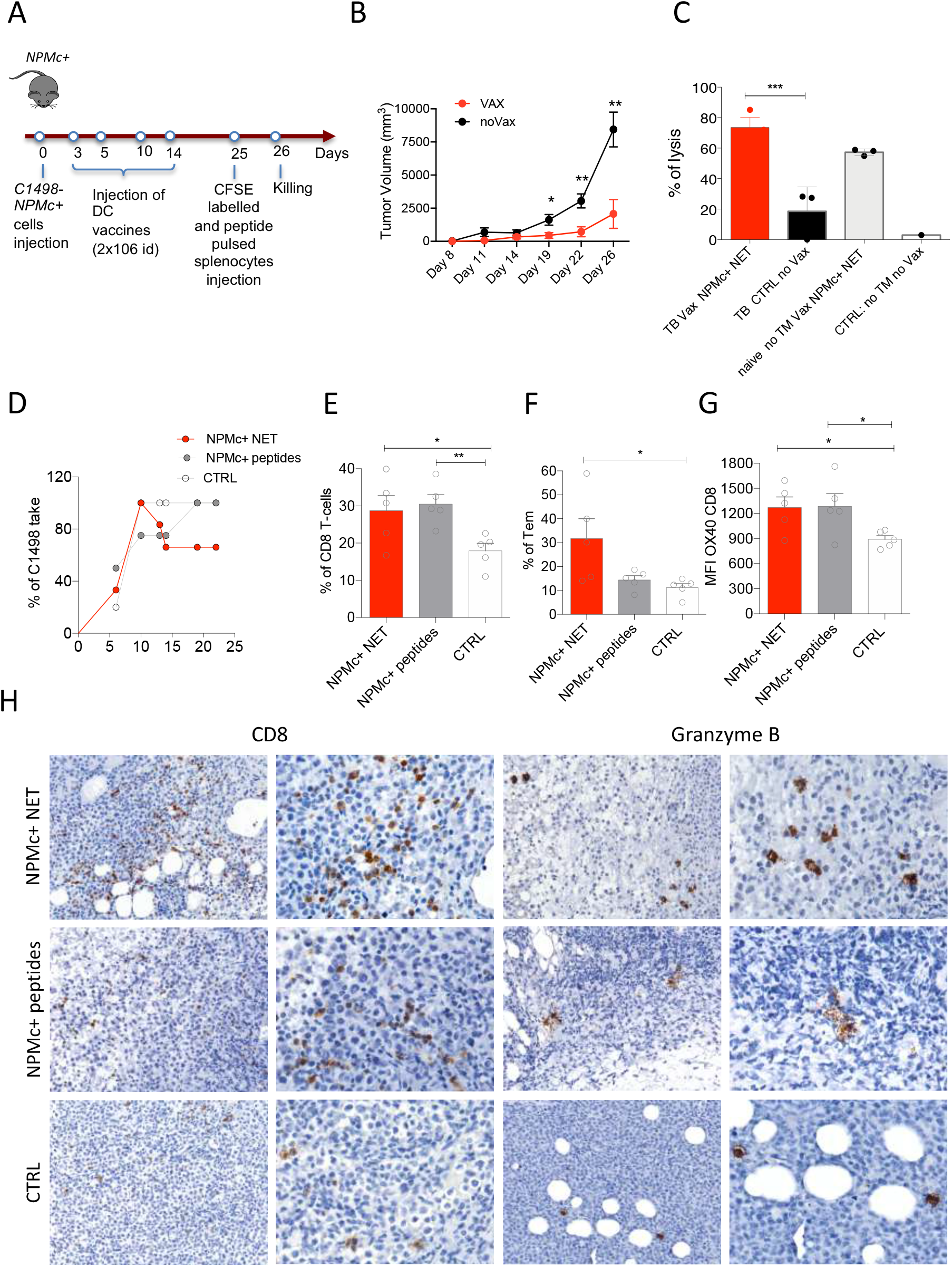
NPMc+ NET vaccination prevents transplantable NPMc+ leukemia cell growth and promotes CD8 lysis. A. Schematic representation of the vaccination experiment; B. C1498-NPMc+ leukemia cells were injected s.c. into NPMc+ transgenic mice. NPMc+ NET/DC-based vaccine was administered at day 3, 5, 10 and 14 post leukemia injection. Tumor growth was monitored twice a week. Vaccination significantly reduced leukemia growth. C. Elicitation of antigen-specific CD8+ T cells in vaccinated mice. Vaccinated tumor bearing mice (TB) or control mice have been injected with 10^7^ cells containing equal numbers of splenocytes labelled with 1.25 µM (CFSE^hi^) or 0.125 µM of CFSE (CFSE^low^). CFSE^hi^ cells were previously pulsed 1 hour with NPMc-MHC-I peptides. Mice were sacrificed the following day, and their splenocytes and lymph nodes analyzed by flow cytometry for the evaluation of the presence of CFSE^hi^ and CFSE^low^ cells. NPMc-specific cytolytic activity was calculated as: (percentage CFSE^high^ cells) × 100/(percentage CFSE^low^ cells). Statistical analysis: one-way ANOVA; Tukey’s multiple comparison test. ***p<0.0001 (n=3/group. One representative experiment out of 3 performed). D. Take of C1498-NPMc+ cells in NPMc+ transgenic mice vaccinated with DC pulsed with NPMc+ NET or NPMc+ peptides at day 3, 5, 10 and 14 post leukemia injection. (n=6/group; Mann-Whitney t test; * p< 0.05, ** p< 0.001). FACS analysis for CD8 T-cell (E), Tem (F) and OX40 expression on CD8 T-cells (MFI)(G). Statistical analysis: one-way ANOVA; Tukey’s multiple comparison test. *p<0.05, **p<0.001 (n=5/group). H. IHC analysis for CD8 and granzyme B of C1498 tumors subcutaneously grown in NPM1 tg mice that received the different vaccinations. (The relative quantification is shown in Suppl. Figure 1B).

The immunization significantly delayed tumor growth (Figure 4B) and induced CD8+ T-cell cytotoxicity against NPMc (Figure 4C). In a different arm of the same experiment, the NPMc*+* NET/DC vaccine also induced a strong CD8+ T cytotoxic response in tumor-free mice (Figure 4C). We also compared the effects of vaccination by DC loaded with NPMc*+* NET and DC loaded with the NPMc*-*derived MHC-I-binding peptides QNYLFGCE, VEAKFINY, and LAVEEVSL. The NPMc*+* NET/DC vaccine proved to be superior in controlling tumor growth (Figure 4D) and in promoting the development of effector-memory CD8+ T-cells (CD8+ Tem). Indeed, both vaccines were able to increase the frequency of total CD8+ T-cells in draining lymph nodes (Figure 4E), but the frequency of CD44+CD62L^neg^ CD8+ Tem was higher in mice vaccinated with NPMc*+* NET/DC than in those receiving DC loaded with NPMc*-*derived peptides (Figure 4F), while the frequency of activated OX40+ CD8+ T-cells was comparable in the two conditions (Figure 4G). In situ IHC analysis of tumor-infiltrating CD8+ T cells and of Granzyme B+ elements confirmed the induction of a cytotoxic response at the site of tumor growth, which was more intense in NPMc*+* NET/DC vaccinated mice in comparison to those vaccinated with NPMc-derived MHC-I peptides or control mice (Figure 4H and Suppl. Figure 1B).

These results collectively demonstrate the actual efficacy of a NPMc*+* NET/DC vaccination approach for the control of an aggressive NPMc+ myeloid malignancy.

## Discussion

Cancer immunotherapy should be able to overcome the induction of drug resistance characterizing standard tumor treatments and to establish effective memory response for durable therapeutic effect independent of repetitive cycles of therapy, with benefits in terms of treatment especially against early lesions or in adjuvant settings. DC loaded *ex vivo* with tumor antigens have largely been tested as cancer vaccines to induce Th1-type responses and trigger cytotoxic T cells targeting antigen-expressing tumor cells.

This strategy has been proposed for patients with AML, with variable clinical benefits (reviewed in (15)). Nevertheless, common weaknesses emerged from different clinical trials including the low ability of DC to mount an effective CD8 T-cell response because of a weak immuno-stimulatory activity and the development of tolerogenic IDO-expressing DC favored by the engendering of an immunosuppressive microenvironment by leukemic clones (16)

This work represents a proof of concept study focusing on NET-based vaccination of NPMc*+* AML. This subset of AML, representing about 30% of total AML and 50-60% of adult normal karyotype-AML, represents a distinct entity in the World Health Organization classification of Tumours of Haematopoietic and Lymphoid Tissues (17). *NPM1*-mutant AML encompasses a spectrum of biological heterogeneity driven by the co-occurrence of genetic mutations (e.g. FLT3 internal tandem duplications) and emerging at the transcriptional level (18).

NPMc+ AML has been characterized for the occurrence of spontaneous NPMc+ NET formation in the BM, which is associated with signs of immune activation ^8^. Despite the pro-immune conditions resulting from a NET-permissive BM environment, the inflammatory skewing and the leukemic clone immunomodulatory profile associated with signs of myelomonocytic differentiation could eventually foster the development of a tolerogenic milieu suitable to NPMc+ AML progression, a condition potentially reverted by vaccination.

The main novelty of our work consists of the adoption of AML blast-derived NET as the source of DC priming. This NET-based approach enables the displaying of leukemia-specific and -associated antigens, such as NPMc, and the conveyance of co-stimulatory signals through the dsDNA thread. In the case of NPMc+ AML, we demonstrated that mutant NPMc exerts a potent adjuvant function, similar to that described for the alarmin HMGB1 (8).

Our results show that NPMc+ NET/DC vaccination is able to break tolerance against NPMc, eventually promoting a strong cytotoxic activity in naive and in AML-bearing NPMc+ Tg mice. These experimental results could represent an important complement to the reported effect of anti-PD-1 immunotherapy in inducing antigen-specific cytotoxic T-cell responses against immunogenic epitopes derived from mutant *NPM1*, which indicates an existing antigen-specific CD8 response in NPMc+ AML patients (19). By comparing NPMc+ NET/DC vaccines with DC pulsed with NPMc-derived peptides, we found a superior capacity of NET-based vaccines to promote anti-tumor immunity inducing CD8 T-cell effector memory responses along with the development of NPMc-specific antibodies. Our results indicate that the adoption of a native form of the mutant protein, such as the one exposed through NETosis, which retains conformational epitopes and adjuvant function, represents a novel promising strategy to achieve immune system awakening against myeloid malignancy-associated antigens.

One potential limitation of the proposed NET/DC vaccination strategy lies in the possibility of the development of autoimmune responses against NET-exposed self antigens, as in the case of anti-neutrophil cytoplasmic antibodies induced against myeloperoxidase or proteinase-3. Autoimmune vasculitides of small vessels have been detected in peripheral tissuess as the result of NET-loaded DC intra-peritoneal injection (6), but no signs of autoimmunity were observed upon adoption of a subcutaneous NET/DC vaccination, as the one used for the vaccination of C1498-NPMc+ tumor-bearing transgenic mice. Another potential caveat is related to the long-term effects of the immune response elicited by NPMc+ NET/DC vaccination on the hematopoietic niche and on its capability to sustain non-malignant hematopoiesis. In our experiments using BM chimeras, we demonstrated that NPMc+ NET/DC vaccination selectively impaired the NPMc+ elements, not disrupting the WT hematopoiesis. However, if the induction of sustained immune responses targeting BM resident elements can in turn favor the establishment of pro-inflammatory conditions contributing to BM niche disfunction is hard to envisage. Indeed, BM hematopoietic stromal niche alterations have been demonstrated to causally affect the arousal and progression of hematopoietic malignancies, either through transcriptional/phenotypical rewiring of key cellular components of the niche (e.g. osteoblasts, nerve fibers, mesenchymal stromal cells) (20–22) or through the modulation of extracellular matrix organization (23). In this regard, extracellular matrix amount and composition have been demonstrated to affect the activation state, cross-presentation activity, and immunoregulatory properties of myeloid elements, an issue that could prove of particular relevance in the unfavorable setting of myeloid malignancies associated with BM fibrosis (24,25).

## Materials and Methods

### Animals

hMPR8-*NPM1* transgenic mice, hereafter referred as *NPM1* tg mice, has been acquired on a mixed (C57BL/6J × CBA) F1 background (8) and backcrossed to C57BL/6 (B6.Ptprc<a>) mice. All experiments involving animals described in this study were approved by the Ministry of Health (authorization number 443/2016-PR e 693/2018-PR).

### Generation of mouse BM chimeras

Competitive bone marrow transplantation assay has been performed transplanting lethally irradiated (1100 RAD) WT mice (B6.Ptprc<a>; CD45.1) with 2×10^5^ Lin negative cells from either *NPMc+* (B6.Ptprc<a>;CD45.1) or WT controls (B6.Ptprc<b>; CD45.2). LK cells were isolated from total BM through negative selection (Lineage Cell Depletion Kit, Miltenyi). This method allows achieving 70% of purity rate.

### Lentiviral vector construction, virus production, cell infection

To construct *NPM1* lentivector the mutant *NPM1* cDNA (mutation type A, TCTG duplication) was cloned into the pLVX-EF1α-IRES-ZsGreen vector (Clontech) using EcoRI and BamHI restriction sites.

A third-generation packaging system was used to produce viral particles. Lentiviral supernantants for *NPM1*-expressing virus and control virus (empty pLVX-EF1α-IRES-ZsGreen vector) were produced in 293T cells by Ca3PO4 co-trasfection of the plasmids as previously described (26). C1498 cell line was infected using viral supernatants at 1:2 ratio with RPMI complete medium. Percentage of infection was evaluated by flow cytometry for GFP expression. Subsequently, infected C1498 cells were sorted by FACSAria to obtain pure populations (100% GFP expression).

### NET-based vaccination protocol

Neutrophils prone to extrude traps have been isolated from agar plugs implanted subcutaneously in WT and *NPM1* transgenic mice, as previously described (6). In particular, PMN were seeded onto coated tissue culture dishes in IMDM 2% FCS, allowed to adhere for 30 minutes and added with myeloid DC (1:1; PMN:DC) for 16h. During this period NET are induced and transfer their component to DC that in turn up-regulate MHC-II and co-stimulatory molecules (6). Then DC are isolated from the co-culture via positive selection, counted and injected (Miltenyi, CD11c Microbeads Ultra Pure). We used 2 schedules for the administration of the DC NET-based vaccines: a) ip injection: 2,5×10^6^ cells, once a week for a total of 6 injection followed by a boost 3 months later; (Schedule in Figure 1A) or b) intradermal: 2×10^6^ cells at day 3, 5, 10 and 14 post leukemia cell injection, Schedule in Figure 4A).

For peptide based vaccination protocol, BM-derived DC have been treated with LPS (10 ng/ml) for 16h and then added with a mixture of 3 *NPMc*+ MHC-I-binding synthetic peptides for 2h. DC were washed and used in vaccination experiments. MHC-I H2-Kb peptides for *NPM1*-A mutant NPMc protein were designed according to the best score interrogating the epitope prediction site syfpeithi.de. Peptides with the following amino acidi sequences, QNYLFGCE, VEAKFINY, LAVEEVSL, were purchased from Primm Biotech (Milan, Italy). Peptides were resuspended in DMSO and then in distilled water.

### Flow cytometry

Staining for cell surface markers was performed in phosphate-buffered saline (PBS) supplemented with 2% fetal bovine serum (FBS) for 30 min on ice. Flow cytometry data were acquired on a LSRFortessa (Becton Dickinson) and analyzed with FlowJo software (version 8.8.6 and 10.4.2, Tree Star Inc.). To assess myeloproliferation in *NPMc*+ transgenic mice, emphasis has been given to the evaluation of the expansion of myeloid precursors. These cells has been identified tanks to their co-expression of CD11b, GR-1 and c-Kit+ markers as described (11). Memory CD4 and CD8 T cell have been identified within their respective gates according to the expression of CD44 and CD62L. Megakaryocyte precursors have been identifiend within the Lin-c-Kit-gate according to their co-expression of CD41 and CD150 markers. All the antibodies that have been used in flow cytometry are listed in Suppl. Table 1.

### In vivo Cytotoxicity Assays

Mice were immunized as described and euthanized one week after the last boost. In vivo cytotoxicity assay was performed as described in (27). Briefly, the day before sacrifice, mice were injected i.p. with 10^7^ cells containing equal numbers of splenocytes labeled with 1.25 mg/ml (CFSE^hi^) or 0.125 mg/ml of CFSE (CFSE^low^). CFSE^hi^ cells were previously pulsed 1h with a mixture of three different NPMc-derived peptides (Peptide sequences: QNYLFGCE; VEAKFINY; LAVEEVSL). Upon sacrifice, splenocytes were analyzed by flow cytometry for the presence of CFSE^hi^ and CFSE^low^ cells. NPMc-specific cytolytic activity was calculated as: (percentage of CFSE^high^ cells) x100 /(percentage of CFSE^low^ cells).

### Histopathological analysis, immunohistochemistry and immunofluorescence

To assess myeloproliferation, the BM of NPMc Tg vaccinated or control mice has been histopathologically evaluated for cellularity, expansion of myeloid cells and degree of maturation of myeloid components. Histopathological analysis was performed on routinely-stained Hematoxylin-and-Eosin sections by a pathologist with specific expertise in hematopathology and murine pathology (CT). Briefly, tissue samples were fixed in 10% buffered formalin, decalcified using an EDTA-based decalcifying solution (MicroDec, Diapath) and paraffin embedded. Four-micrometers-thick sections were deparaffinized and rehydrated. The antigen unmasking technique was performed using Novocastra Epitope Retrieval Solutions pH6 and pH9 in a thermostatic bath at 98°C for 30 minutes. Subsequently, the sections were brought to room temperature and washed in PBS. After neutralization of the endogenous peroxidase with 3% H_2_O_2_ and Fc blocking by a specific protein block (Novocastra UK), the samples were incubated with the primary antibodies. The following primary antibodies were adopted for IHC and IF: NPM (clone 376, dilution 1:100 pH6, Dako); CD8α (clone D4W2Z, dilution 1:400 pH9, Cell Signaling) and Granzyme B (dilution 1:10 pH6, Cell Marque). The staining was revealed using IgG (H&L) specific secondary antibody HRP-conjugated (1:500, Novex by Life Technologies) and DAB (3,3’-Diaminobenzidine, Novocastra) as substrate chromogen. For double-marker immunofluorescence, after antigen retrieval, the sections were incubated overnight at 4°C with NPM and CD8α primary antibodies. The binding of the primary antibodies to their respective antigenic substrates was revealed by made-specific secondary antibodies conjugated with Alexa-488 (Life Technologies, 1:250) and Alexa-568 (Life Technologies, 1:300) fluorochromes. Nuclei were counterstained with DAPI (4’,6-diamidin-2-fenilindolo). Slides were analyzed under an Axioscope A1 microscope equipped with an Axiocam 503 Color digital camera and Zen 2.0 Software (Zeiss). Quantitative IHC data were obtained by calculating the number of CD8^+^ or Granzyme B^+^ cells in five non-overlapping fields at high-power magnification (x400).

To measure the cell-cell contact between NPMc+ cells (green signal) and CD8+ cells (red signal), we applied the redundant wavelet transform called à trous (28) (29) because it preserves the original resolution of the image while allowing the removal of noise. With such a method, the quality of the data was preserved without the need of pre or post-processing techniques. The intensity of overlay signal (yellow signal) was then measured only within the significant zones identified automatically and related to their own areas. This approach makes the methodology we developed independent of the zoom factor used to acquire the photomicrographs.

### Confocal microscopy analysis

To evaluate NET formation, NPMc re-localization onto the NET threads, and the transfer of MPO and NPMc+ antigens to DC, agar neutrophils from WT and NPMc*+* mice were seeded onto poly-D-lysine coated glasses (IMDM 2% FCS), allowed to adhere for 30 minutes and added with mDCs (1:1 PMN:DC). After 16h, cells were fixed in 4% PFA and sequentially stained with mAbs to NPM (anti-human/mouse nucleophosmin (NPM) (clone 376, dilution 1:50, Dako) and anti-mouse MPO (Millipore). The NET-DNA was counterstained with Draq5 or the vital DNA dye Sytox green, or with Dapi. IF stainings were acquired under a Leica TCS SP8 confocal microscope (Leica Microsystems).

### Statistical Analysis

All statistical analyses have been performed with Prism 8 (GraphPad Software). The statistic applied to every single experiment is shown in the relative figure legend. We applied both parametric and non-parametric analysis including Student t test, Mann-Whitney test according to data distribution. In case of multiple comparison we used the one-way ANOVA analysis with Tukey’s or Dunnett’s multiple comparison.

## Acknowledgements

This work has been supported by the Italian Ministry of Health (GR-2013-02355637 to S. Sangaletti) and Italian Cancer Research Foundation (AIRC) (Investigator Grant number 22204 to S. Sangaletti). The authors are grateful to Ester Grande for administrative support.

## Supplemental Material

**Supplementary Figure 1.**
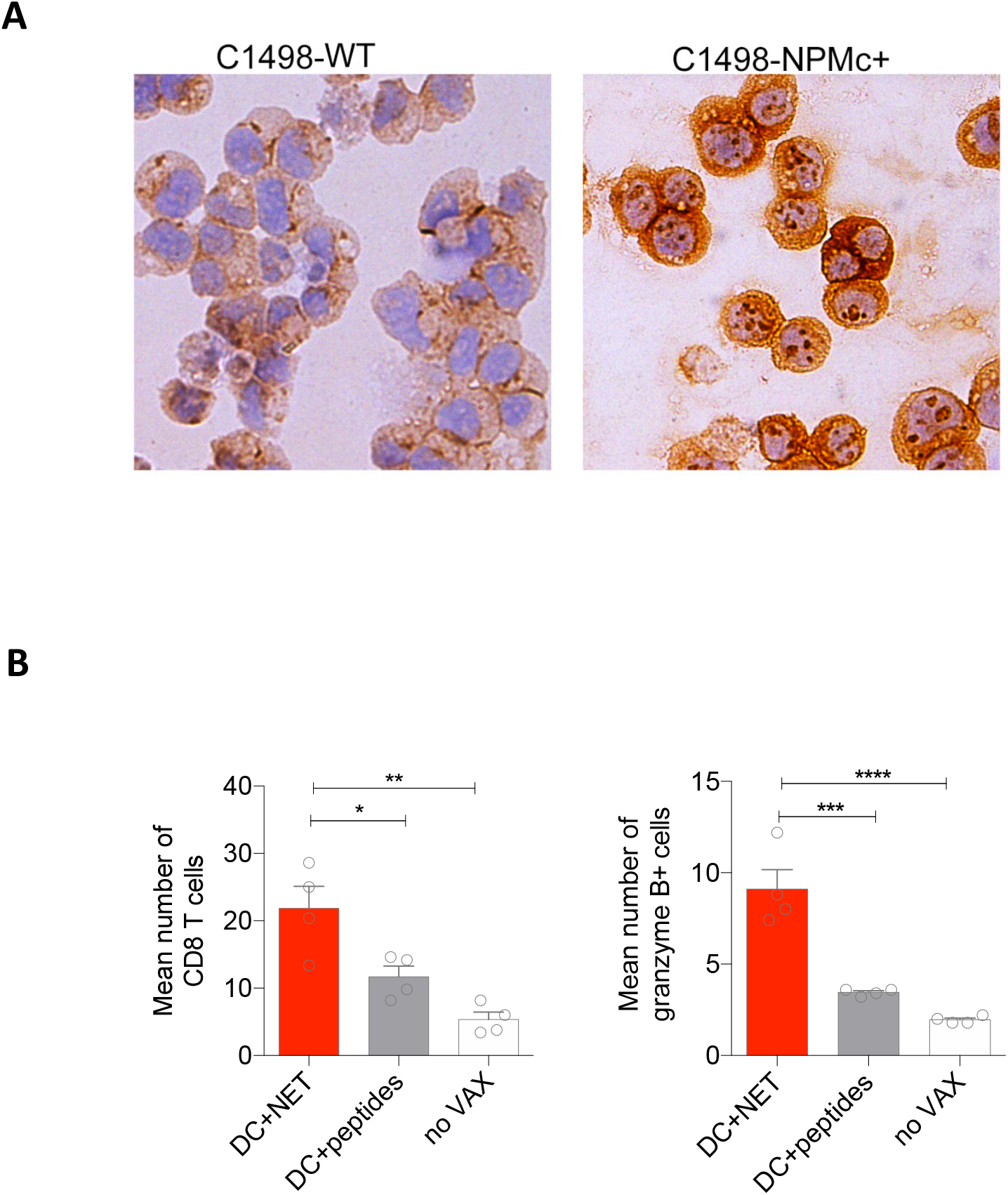
A. IHC analyses for mutant NPMc in parental vs lentiviral infected C1498 cells; B. Quantification of the mean number of CD8 T cell and of granzyme B+ cell in C1498 tumors obtained from mive vaccinated with DC+NET or DC+peptides. Statistical analysis: one-way ANOVA, Tukey’s multiple comparison test. * p< 0.05; *** p< 0.001.

**Supplementary Table 1.**
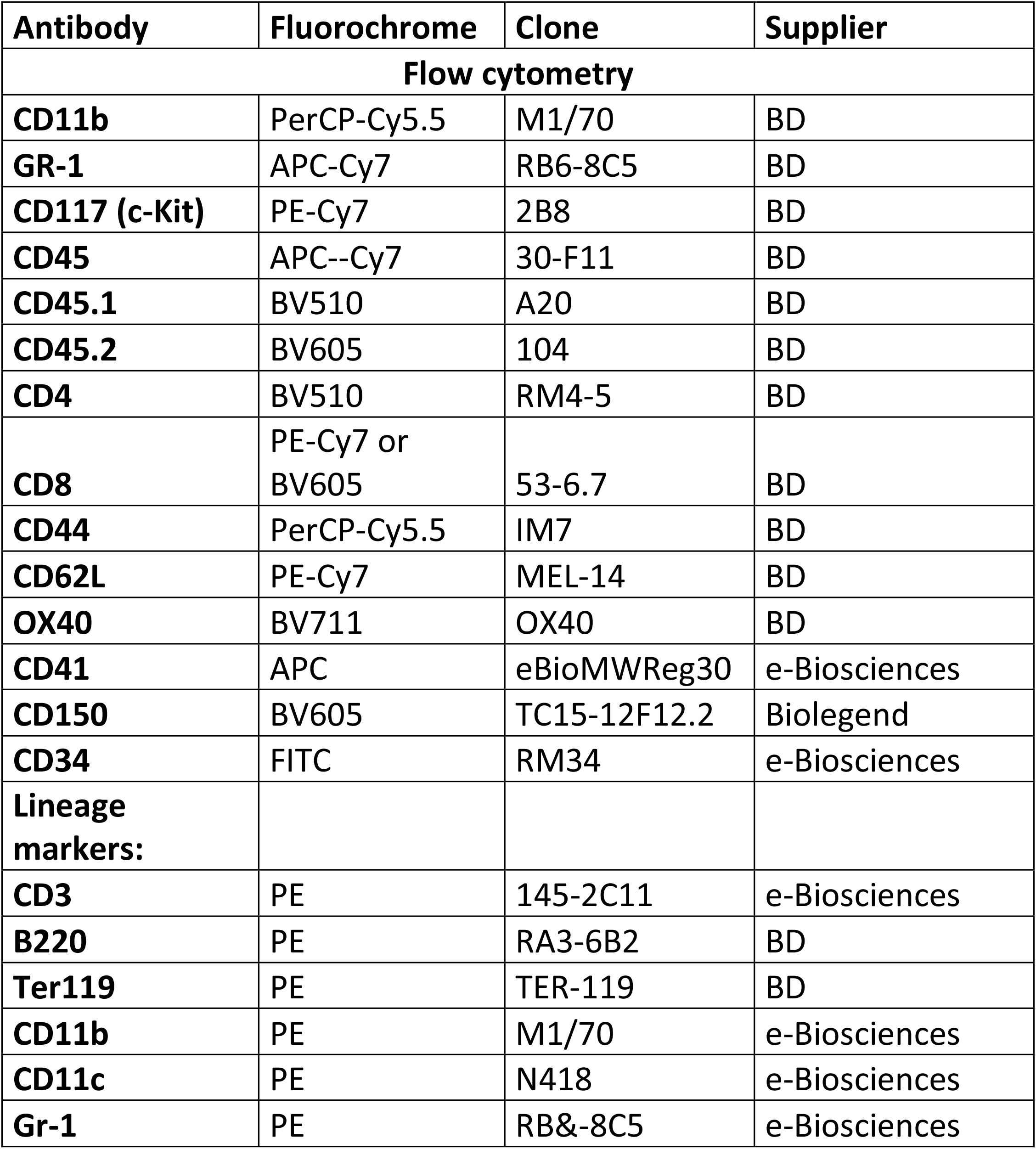
Antibodies for flow cytometry

## Notes

### Competing Interest Statement

The authors have declared no competing interest.

